# System-wide biochemical analysis reveals ozonide antimalarials initially act by disrupting *Plasmodium falciparum* haemoglobin digestion

**DOI:** 10.1101/2020.03.23.003376

**Authors:** Carlo Giannangelo, Ghizal Siddiqui, Amanda De Paoli, Bethany M. Anderson, Laura E. Edgington-Mitchell, Susan A. Charman, Darren J. Creek

## Abstract

Ozonide antimalarials, OZ277 (arterolane) and OZ439 (artefenomel), are synthetic peroxide-based antimalarials with potent activity against the deadliest malaria parasite, *Plasmodium falciparum*. Here we used a “multi-omics” workflow, in combination with activity-based protein profiling (ABPP), to demonstrate that peroxide antimalarials initially target the haemoglobin (Hb) digestion pathway to kill malaria parasites.

Time-dependent metabolomic profiling of ozonide-treated *P. falciparum* infected red blood cells revealed a rapid depletion of short Hb-derived peptides followed by subsequent alterations in lipid and nucleotide metabolism, while untargeted peptidomics showed accumulation of longer Hb-derived peptides. Quantitative proteomics and ABPP assays demonstrated that Hb-digesting proteases were increased in abundance and activity following treatment, respectively. The association between ozonide activity and Hb catabolism was also confirmed in a *K13*-mutant artemisinin resistant parasite line. To demonstrate that compromised Hb catabolism may be a primary mechanism involved in ozonide antimalarial activity, we showed that parasites forced to rely solely on Hb digestion for amino acids became hypersensitive to short ozonide exposures.

Quantitative proteomics analysis also revealed parasite proteins involved in translation and the ubiquitin-proteasome system were enriched following drug treatment, suggestive of the parasite engaging a stress response to mitigate ozonide-induced damage. Taken together, these data point to a mechanism of action involving initial impairment of Hb catabolism, and indicate that the parasite regulates protein turnover to manage ozonide-induced damage.

**Author Summary:** The ozonides are a novel class of fully synthetic antimalarial drugs with potent activity against all parasite species that cause malaria, including the deadliest, *Plasmodium falciparum*. With the emergence of resistance to current frontline artemisinin-based antimalarials, new drugs are urgently needed and a clear understanding of their mechanism of action is essential so that they can be optimally deployed in the field. Here, we studied the biochemical effects of two ozonides, OZ277 (marketed in India in combination with piperaquine) and OZ439 (in Phase IIb clinical trials) in *P. falciparum* parasites using an untargeted multi-omics approach consisting of proteomics, peptidomics and time-dependent metabolomics, along with activity-based protease profiling. We found that the ozonides initially disrupt haemoglobin metabolism and that they likely engage the parasite proteostatic stress response. Furthermore, when the duration of ozonide exposure was extended beyond 3 hours to reflect clinically-relevant exposure periods, additional parasite biochemical pathways were perturbed. This comprehensive analysis provides new insight into the antimalarial mode of action of ozonides and provides new opportunities for interventions to enhance their antimalarial efficacy.

## Introduction

Promising improvements in malaria control have been recorded over the last two decades, but recent data indicates that the declining mortality rates have either stalled or increased in many malaria endemic regions since 2016 [1]. The absence of a reliable and highly efficacious vaccine means that treatment is heavily reliant on effective antimalarial chemotherapy. Currently, the World Health Organisation (WHO) recommends artemisinin-based combination therapies (ACTs) as the first-line treatment for uncomplicated *Plasmodium falciparum* malaria in all endemic areas [2]. The artemisinins (including dihydroartemisinin, DHA) contain an essential peroxide bond that undergoes reductive activation by haem released through parasite haemoglobin (Hb) digestion [3–6]. This activation process generates highly reactive drug-derived radicals that mediate rapid parasite killing [7]. However, artemisinins are limited by poor biopharmaceutical properties and short *in vivo* half-lives (< 1 h) [7–9]. Furthermore, the emergence of artemisinin resistant parasites now threatens global malaria control and elimination efforts [10]. Thus, there is a desperate need for improved therapeutics to combat malaria.

To overcome some of these limitations, the artemisinin peroxide bond inspired the design of fully synthetic and structurally dissimilar peroxide-based antimalarials, known as ozonides [11] (Fig. 1). The first-generation ozonide, OZ277 (later known as RBx11160 or arterolane) [11], was the first to be approved clinically and is currently marketed as a fixed dose combination with piperaquine (Synriam^TM^). However, the *in vivo* half-life of OZ277 is only 2- to 3-fold longer than that for DHA [12, 13]. This rapid clearance is thought to be associated with both hepatic metabolism [14] and instability of the peroxide bond when exposed to endogenous sources of iron in blood and tissues [15]. A design strategy aimed at stabilising the peroxide bond to iron-mediated degradation led to the development and selection of the second-generation ozonide, OZ439 (artefenomel) [15]. When tested clinically, OZ439 exhibited an *in vivo* half-life of 46-62 h in humans [16, 17]. OZ439 is currently in Phase IIb clinical trials in combination with ferroquine (ClinicalTrials.gov Identifier: NCT02497612).

**Fig. 1.**
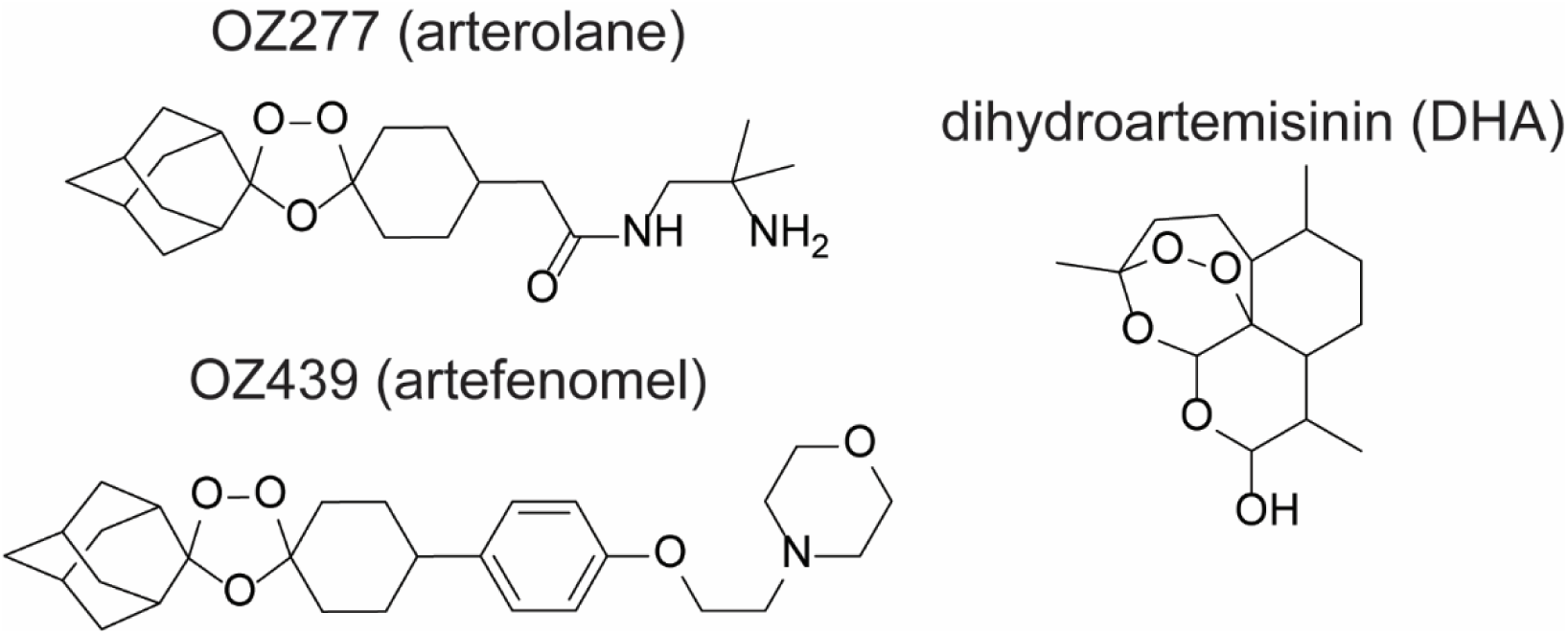
Chemical structures of selected peroxide antimalarials. The fully synthetic ozonide antimalarials, OZ277 (arterolane) and OZ439 (artefenomel), and the clinically used semisynthetic artemisinin derivative, dihydroartemisinin (DHA).

Ozonide antimalarials display similar clinical efficacy to DHA, rapidly clearing blood-stage parasites [17, 18]. However, their antimalarial mechanism of action (MoA) has been less extensively studied than the artemisinins, and debate remains about the key molecular events responsible for artemisinin action [19–24]. The current model for ozonide antimalarial activity is that Hb-derived free haem mediates reductive activation of the peroxide bond giving rise to toxic carbon-centred radicals [25] that alkylate a number of essential parasite proteins from various biochemical pathways [26, 27]. Crucially, optimal clinical utilisation of ozonides will rely on a clear understanding of the biochemical mechanisms that underpin their activity. Therefore, we investigated the temporal biochemical response of *P. falciparum* parasites treated with OZ277 and OZ439 using systems-wide analyses, incorporating time-dependent metabolomics and proteomics. We identified Hb digestion as the key initial pathway targeted by ozonide antimalarials and demonstrated that when parasites are forced to rely solely on Hb digestion for nutrients, they become hypersensitive to pulsed ozonide treatment. Furthermore, we showed that ozonides perturb additional pathways when treatment was extended beyond 3 h, reflecting more clinically-relevant exposures for the ozonides, and that parasites likely regulate protein turnover to manage ozonide-mediated damage. This work provides new opportunities for interventions to target malaria parasites and enhance ozonide antimalarial efficacy.

## Results

### Ozonide antimalarials initially deplete short haemoglobin (Hb)-derived peptides

In order to distinguish the early peroxide-induced effects from secondary mechanisms, we employed a time-dependent, untargeted metabolomics approach that allowed comprehensive biochemical profiling of the primary pathways affected by ozonide antimalarials in *P. falciparum* infected red blood cells (iRBCs) (Supplementary Fig. 1). Trophozoite-stage parasites (28-34 h post invasion) were treated with 300 nM of OZ277 or OZ439 or 100 nM of DHA over a time course of up to 3 h (n = at least four biological replicates). These drug concentrations are equivalent to the IC_50_ for a 3 h pulse, under the same *in vitro* conditions used in the metabolomics analysis (10% parasitaemia and 2% Hct) [28], and are within therapeutic concentration ranges [10, 17, 29]. Univariate analysis of the untargeted metabolomics dataset (Supplementary Dataset 1) revealed a temporal increase in the percentage of drug-induced metabolic changes (Supplementary Fig. 2a) and that widespread metabolic perturbations were not evident after short drug exposures (Supplementary Fig. 2b). The ozonides rapidly and disproportionately affected peptide metabolism, with approximately 15-25% of all putatively identified short peptides (2-4 amino acids in length) significantly perturbed within 3 h of drug treatment (P-value < 0.05) (Supplementary Fig. 2b). These significantly perturbed peptides were parasite-specific metabolites (Supplementary Dataset 1) and exhibited a progressive depletion in abundance over the 3 h drug exposure (Fig. 2a and 2b). Interestingly, the extent of peptide depletion was more extensive and faster in DHA-treated parasites compared to the ozonides, which is consistent with the more rapid kinetics of antiparasitic activity previously observed for DHA compared to the ozonides for these short pulse exposures (< 3 h) [30]. The amino acid sequence of a subset of these putatively identified peptides was confirmed by MS/MS and the majority of perturbed peptides (i.e., those that were decreased by at least 1.5-fold) could be mapped to either the alpha or beta chains of Hb (Fig. 3a).

**Fig. 2.**
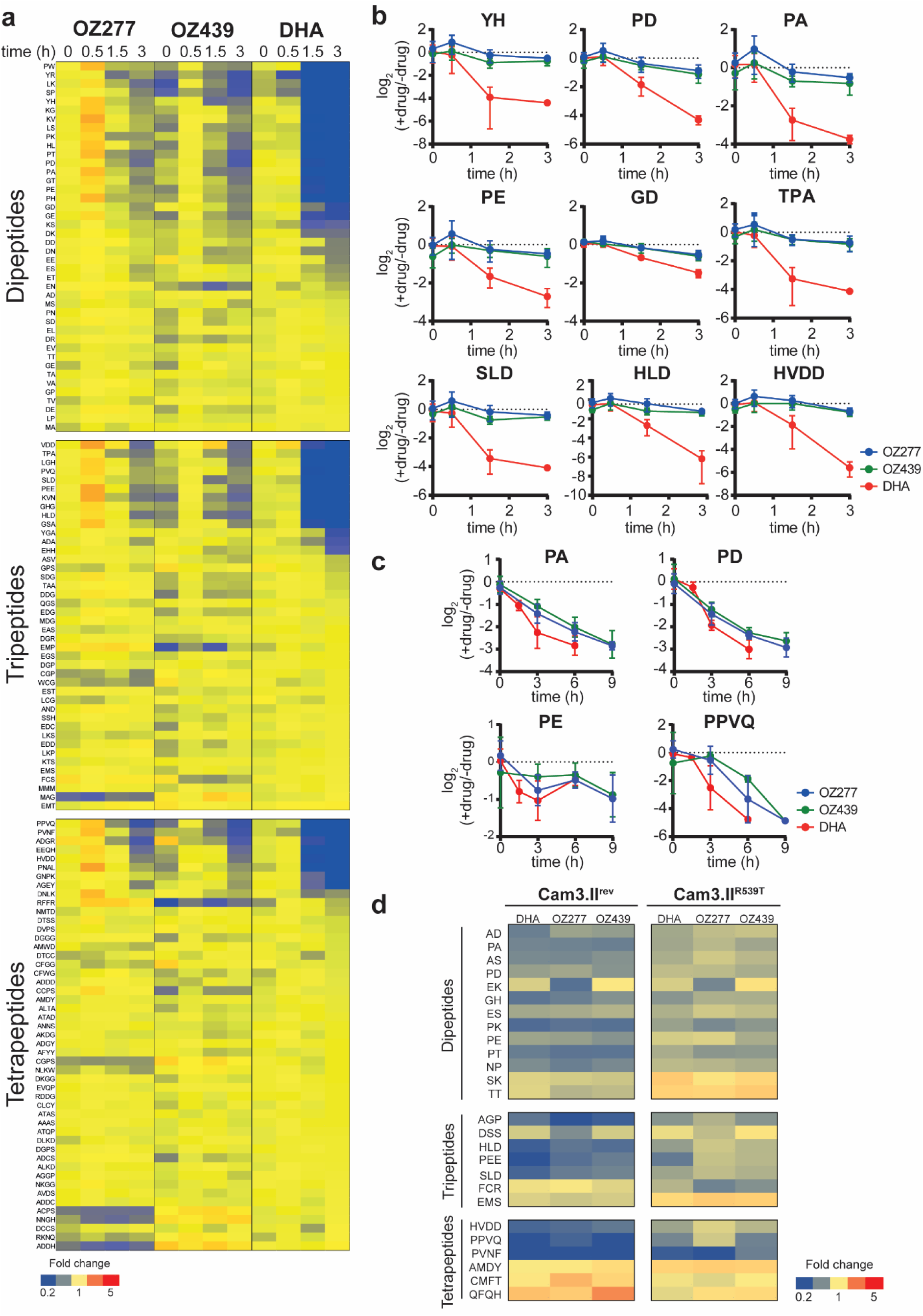
Peroxide-induced perturbations to peptide metabolism. **a**, Heatmap showing the average fold change for all identified peptides at each time point after treatment with OZ277, OZ439 and DHA. Values represent the average of at least three biological replicates, expressed relative to the average untreated control value (at least seven biological replicates) for that respective time point. **b**, Representative time profiles showing the progressive depletion in abundance of selected putative haemoglobin-derived peptides after peroxide treatment of trophozoite-stage parasites. Values are the average fold change (± SD) relative to the untreated control of at least three biological replicates. **c**, Time-dependent decrease in the abundance of the four peptides consistently depleted following peroxide treatment in ring infected cultures. Values are the average fold change (± SD) relative to the untreated control of four biological replicates. **d**, Heatmap of the peptides that were altered in abundance (≥ 1.5-fold relative to the untreated control) following peroxide treatment in the *K13*-wildtype artemisinin sensitive (Cam3.II^rev^) and *K13*-mutant artemisinin resistant (Cam3.II^R539T^) parasite lines. Data shown are the average for three or five biological replicates expressed relative to the average for the untreated control from the same parasite line.

**Fig. 3.**
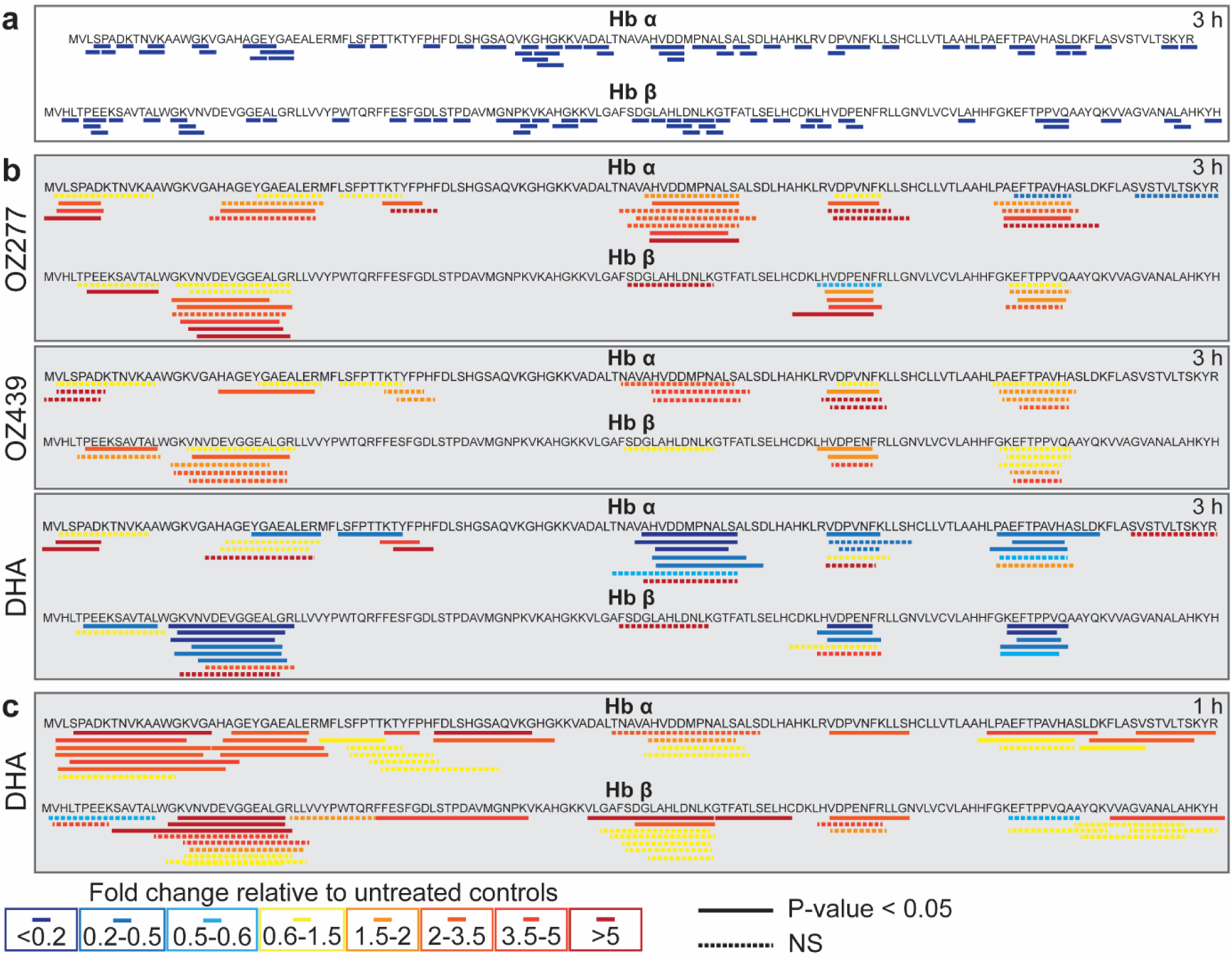
Sequence coverage and relative abundance of haemoglobin alpha (Hb α) and haemoglobin beta (Hb β) peptides detected after peroxide treatment. **a**, Sequence coverage for putative Hb-derived di-, tri-, and tetra-peptides that were differentially abundant (≥ 1.5-fold) following peroxide antimalarial treatment relative to the untreated control (3D7 parasites). The peptide sequences PA, PT, PE, PEE, HLD, SLD, PPVQ, PVNF and HVDD have been confirmed by MS/MS analysis. For all other putative peptide sequences, all potential isomers have been mapped. **b**, Sequence coverage for all long Hb α and Hb β peptides detected in peptidomics studies. Peptide abundances are the average fold change following 3 hours of drug treatment (OZ277, OZ439 and DHA), expressed relative to the untreated control (DMSO) from at least three biological replicates (3D7 parasites). **c**, Sequence coverage for Hb α and Hb β peptides detected following 1 hour of DHA treatment in Cam3.II^rev^ (artemisinin sensitive) parasites. Peptide abundances are the average fold change expressed relative to the untreated control (DMSO) from two biological replicates. In **b**, **c**, samples were normalised according to peptide concentration (measured using a bicinchoninic acid assay) during sample preparation. The solid lines represent the amino acid sequences of peptides that significantly changed in abundance after drug treatment relative to the untreated control (P-value < 0.05). Dashed lines represent non-significant (NS) changes. Increased and decreased peptide abundance are represented by red and blue (solid or dashed) lines, respectively.

A similar, although less extensive, temporal increase in the number of significant metabolic perturbations was confirmed in ring-stage parasites (6-12 h post invasion) treated with ozonide antimalarials (Supplementary Fig. 3a), with putative Hb-derived peptides representing the first metabolites to be significantly perturbed (Supplementary Dataset 1 and Supplementary Fig. 3b). This was further verified using multivariate analysis, where sparse partial least squares– discriminant analysis (sPLS-DA) revealed that Hb-derived short peptides were responsible for the greatest differences between the ozonide-treated samples and controls (Supplementary Fig. 4). Similar to trophozoite-stage parasites, these Hb peptides showed a time-dependent depletion in the treated parasite cultures compared to controls (Fig. 3c).

To further investigate the importance of parasite Hb digestion to ozonide antimalarial action, untargeted metabolomic profiling was also performed on artemisinin resistant (Cam3.II^R539T^) and isogenic sensitive (Cam3.II^rev^) [31] early trophozoite-stage (22-26 h post invasion) parasites (Supplementary Dataset 2). Artemisinin resistant early trophozoite-stage parasites have been shown to exhibit differential sensitivity to short DHA exposures compared to the isogenic sensitive strain, albeit a less dramatic difference than early rings [32]. Targeted analysis of the LC-MS raw data identified 26 perturbed peptides (at least 1.5-fold) in drug-treated resistant (Cam3.II^R539T^) or sensitive parasites compared to control (Fig. 2d). The majority (> 70%) of perturbed peptides were depleted in treated samples compared to the control (Fig. 2d and Supplementary Fig. 5a) and most of these peptides (all except two) could be mapped to Hb (Supplementary Fig. 5b). Notably, the extent of peptide depletion was greater in the sensitive parasites than in the resistant line (Fig. 2d). In general, the abundance of these peptides in treated resistant parasites remained at, or above, the basal levels detected in untreated sensitive parasites.

### Ozonide antimalarial treatment causes accumulation of long chain Hb peptides

The rapid depletion of short chain Hb-derived peptides led us to consider how ozonide antimalarial exposure impacts longer Hb-derived peptides. We used a MS/MS-based global peptidomics approach (Supplementary Fig. 1) to examine the abundance of endogenous peptides (< 10 kDa) within ozonide-treated *P. falciparum* parasites. Peptidomics analysis identified a total of 59 endogenous *P. falciparum* peptides and 59 endogenous human peptides (Supplementary Dataset 3). OZ277 (300 nM for 3 h) treatment significantly altered the abundance of 30 endogenous peptides in trophozoites-stage parasites (P-value < 0.05), 17 of which originated from Hb (alpha and beta), and were increased in abundance (Fig. 3b). A similar build-up of long chain Hb peptides was also observed following treatment with OZ439 (300 nM for 3 h). Unlike the ozonides, exposure of trophozoite-stage parasites to 3 h of DHA (100 nM) predominantly depleted the abundance of Hb-derived peptides, with 26 peptides significantly depleted and four significantly elevated relative to control (Fig. 3b). The differential impact of ozonides and DHA on longer chain Hb-derived peptides may be explained by the faster onset of action of artemisinins compared to ozonides [30]. Indeed, a shorter DHA exposure (1 h) caused an accumulation of longer Hb peptides similar to the ozonides (Fig. 3c and Supplementary Dataset 4).

We also assessed whether the accumulated Hb components in peroxide-treated parasites differ from those in E64d (cysteine protease inhibitor)-treated parasites. E64d causes parasite digestive vacuoles to accumulate undegraded Hb and swell due to disruption of the initial endoproteolytic cleavage of Hb [33, 34]. Trophozoite-stage parasites treated with E64d for up to 3 h developed a characteristic swollen digestive vacuole (Supplementary Fig. 6a), consistent with abrogated digestion of full-length Hb. These same parasites exhibited a modest trend towards accumulation of intact Hb and minor changes in free haem and haemozoin levels (Supplementary Fig. 6a) when these haem-containing species were measured using haem fractionation assays [35]. Visualisation of monomer Hb (17 kDa) by Coomassie staining of SDS-PAGE gels confirmed that undigested Hb accumulated in trophozoites after these short E64d exposures (Supplementary Fig. 6b). Conversely, peroxide-treated parasites showed a minor decrease in the levels of full-length Hb, when measured using the haem fractionation assay (no changes in other haem species were evident) (Supplementary Fig. 6a), which is broadly consistent with our quantitative proteomics data (Supplementary Fig. 6c). Peroxide exposures of up to 3 h caused no changes in digestive vacuole morphology, indicating that there is no inhibition of proteolysis of full-length Hb (Supplementary Fig. 6a). Taken together, these findings suggest that the accumulated Hb components in peroxide-treated parasites are likely different from those resulting from specific inhibition of cysteine proteases.

Our untargeted peptidomics analysis also showed that ozonide treatment perturbed the levels of some endogenous parasite peptides. Of the 13 parasite peptides significantly perturbed following OZ277 exposure, five originated from an uncharacterised *P. falciparum* protein (PF3D7_0716300) and were decreased in abundance compared to control (Supplementary Dataset 3). OZ439 treatment resulted in significant perturbations to five parasite peptides, four of which were from this same uncharacterised protein (PF3D7_0716300) and were decreased relative to the untreated (Supplementary Dataset 3). A total of 15 endogenous parasite peptides were altered in abundance following DHA treatment, including six peptides that originated from the uncharacterised protein PF3D7_0716300 and, similar to ozonide exposure, were significantly decreased compared to control (Supplementary Dataset 3).

### Ozonide antimalarial treatment increases the abundance and activity of Hb proteases

*Plasmodium* parasites digest Hb through a semi-ordered proteolytic process incorporating numerous proteases of different classes [37]. Peptidomics and metabolomics analyses of treated parasites suggested that ozonide antimalarials disrupt Hb catabolism through inhibition of the proteases involved in the breakdown of large to small Hb peptides. In order to quantify the abundance of the proteases involved in Hb digestion, we used dimethyl labelling-based quantitative proteomics (Supplementary Fig. 1) [38]. Targeted analysis of the global proteomics data (Supplementary Fig. 7 and Supplementary Dataset 5) identified falcipains 2 and 3 (FP 2 and FP 3) and the plasmepsins (PM I, PM II, PM IV and HAP), proteases thought to be involved in the initial stages of Hb degradation [37], to be elevated in abundance in drug-treated samples compared to control (Fig. 4a). Dipeptidyl aminopeptidase 1 (DPAP1), which removes dipeptides from the polypeptides produced by upstream proteases [39], and the alanyl aminopeptidase (*Pf*A-M1), leucyl aminopeptidase (*Pf*A-M17) and aspartyl aminopeptidase (*Pf*M18AAP) metalloproteases, all of which are involved in the terminal stages of Hb digestion [40], were all increased following ozonide treatment (Fig. 4a). No changes in the abundance of falcilysin (Fig. 4a) was detected (Supplementary Dataset 5).

**Fig. 4.**
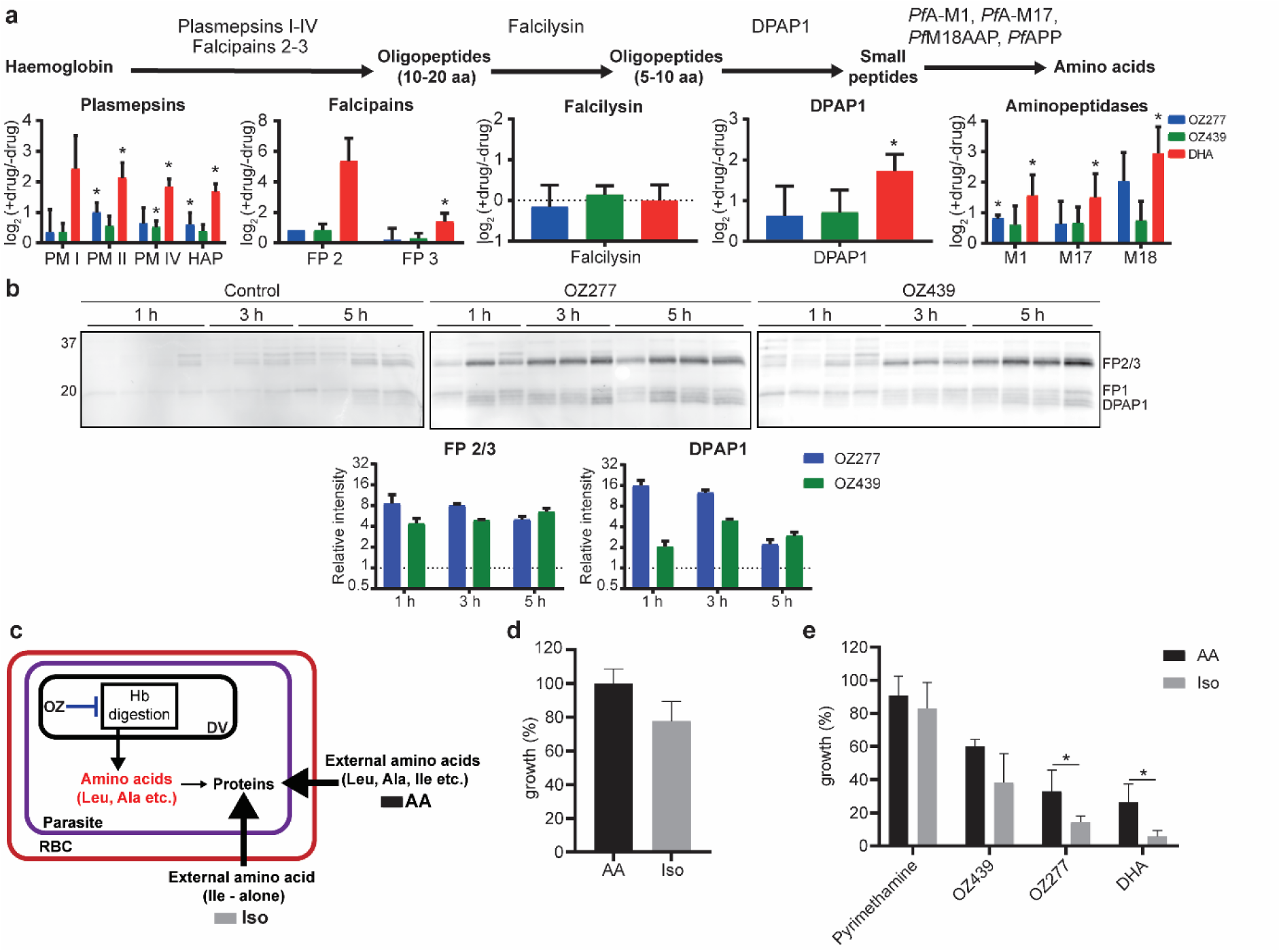
Peroxide antimalarials act by perturbing haemoglobin digestion. **a**, Disruption of protease abundance in the haemoglobin digestion pathway [37]. Values are the average log_2_ fold change (± SD) relative to the untreated control of at least three biological replicates. Trophozoite infected cultures were incubated with OZ277, OZ439 (both 300 nM) or DHA (100 nM) for 3 h. Falcipain 2 (FP 2) was identified in only two OZ277 treatment experiments, therefore the mean alone is shown. Aminopeptidase P (*Pf*APP) was not identified in any of the proteomic experiments. DPAP1, dipeptidyl aminopeptidase 1; FP, falcipain; HAP, histo-aspartic protease; *Pf*APP, aminopeptidase P; *Pf*A-M1, alanyl aminopeptidase; *Pf*A-M17, leucyl aminopeptidase; *Pf*M18APP, aspartyl aminopeptidase; PM, plasmepsin; * P-value < 0.05. **b**, Parasite cysteine protease activity after peroxide treatment using the activity-based probe (ABP), DCG04. Cysteine protease activity and densitometric analysis of the falcipain (FP) 2/3 and dipeptidyl aminopeptidase 1 (DPAP1) signal after OZ277 and OZ439 treatment in *P. falciparum* trophozoite stage parasites using DCG04 in lysates at pH 5.5 (acidic). Trophozoite infected cultures were incubated with OZ277, OZ439 (both 300 nM) or an equivalent volume of DMSO (control). DCG04 labelling was detected by blotting membranes with streptavidin-AF647 after SDS-PAGE and transfer. The lanes for each time point are independent drug treatments and represent at least three biological replicates per time point that were run on the same gel side-by-side. For the densitometric analysis, the post drug treatment FP 2/3 and DPAP1 signal intensity was normalised to the average signal intensity of the appropriate time point in the untreated control (±SD). **c**, Schematic showing that infected RBCs treated with DHA or ozonides (OZ) can use exogenous amino acids when cultured in AA medium (Full RPMI medium with all 20 amino acids) in response to disrupted haemoglobin digestion (arrow shown in blue), while parasites in Iso medium (supplemented with isoleucine alone at a final concentration of 147.5 µM) must rely solely on haemoglobin digestion for amino acids. **d**, Amino acid requirement for cultured *P. falciparum* 3D7 parasites. Parasite viability measured following 48 h incubation in medium containing all amino acids (AA, black bars) and isoleucine alone medium (Iso, grey bars). **e**, Parasite sensitivity to peroxides when cultured in AA (black bars) medium compared to Iso (grey bars) medium. Trophozoite infected cultures were incubated with pyrimethamine, OZ277, OZ439 (all 300 nM), DHA (100 nM) or an equivalent volume of DMSO (control) for 3 h. Data represents the mean ± SD of at least three biological replicates. Growth values for each treatment is expressed relative to the appropriate untreated medium (Iso or AA) control, which was set to 100%. * P-value < 0.05.

As most Hb-degrading proteases were elevated in abundance after ozonide antimalarial treatment, we then investigated the temporal impact of ozonides on Hb protease activity using activity-based probes (ABPs) targeting parasite cysteine proteases [41] (Supplementary Fig. 1). Trophozoite-stage parasite cultures were treated with OZ277 or OZ439 (300 nM) for up to 5 h and the biotinylated epoxide ABP, DCG04 [42], was used to label the Hb-digesting cysteine proteases FP 2, FP 3 and DPAP1 in the parasite lysate under both acidic (pH 5.5) and neutral pH (pH 7.2) conditions. Both OZ277 and OZ439 caused a time-dependent increase in the activity of proteases with molecular weights consistent with that of the FPs (FP 2 and FP 3), and DPAP1 [43] (Fig. 4b). A similar increase in the activity of these proteases was observed under acidic (the pH environment of the parasite digestive vacuole) or neutral pH conditions (Supplementary Fig. 8a), and activity was inhibited by pre-treatment of the parasite lysate with the cysteine protease inhibitor, ALLN (Supplementary Fig. 8b). FPs and DPAP1 activity were increased within 1 h of OZ277 treatment compared with DMSO controls (Fig. 4b). In contrast, OZ439 increased FPs and DPAP1 activity after 3-5 h of drug exposure, consistent with it having a slower onset of action within the parasite [30]. Ozonide-induced increases in activity of FPs and DPAP1 were further confirmed by another cysteine protease-targeting probe, FY01 [43], under both acidic (Supplementary Fig. 9a) and neutral (Supplementary Fig. 9b) conditions.

### Impaired Hb digestion underpins initial ozonide-induced toxicity

Functional Hb uptake and digestion is essential for parasite survival as it supplies amino acids for protein synthesis [44]. As our multi-omics and ABPP analyses identified Hb catabolism as the primary pathway affected by ozonide antimalarial treatment, we hypothesised that drug-derived radicals initially target this pathway, disrupting Hb catabolism, and starving the parasite of Hb-derived amino acids. To test this, we determined the potency of peroxide antimalarials on parasites grown in full RPMI medium (with all 20 amino acids) (AA medium) and parasites cultured in medium lacking all exogenous amino acids except for isoleucine (the only amino acid absent from Hb) (Iso medium), thereby forcing parasites to rely solely on Hb catabolism for amino acid supply (Fig. 4c).

Similar to published results [44], parasites cultured in Iso medium had a minor growth defect of approximately 20% compared to parasites cultured in AA medium (Fig. 4d). Trophozoite-stage cultures exposed to OZ277 or OZ439 (both 300 nM) for 3 h were sensitised by 2.3- and 1.6-fold, respectively, in the Iso medium (Fig. 4e), suggesting that compromised Hb catabolism is the primary MoA of ozonides in the initial exposure phase. However, when drug pressure was maintained throughout the entire RBC life cycle, there was no difference in sensitivity between parasites cultured in the Iso and AA mediums (Supplementary Fig. 10), suggesting that mechanisms beyond disrupted Hb digestion contribute to parasiticidal activity during prolonged exposure. DHA treated parasites were almost 5-fold more sensitive in the Iso medium compared to parasites cultured in AA medium (Fig. 4e). In contrast, the potency of pyrimethamine, which kills parasites by a mechanism independent of Hb digestion [45], was not affected when cultured in either AA or Iso medium (Fig. 4e).

### Ozonide antimalarial treatment upregulates parasite proteasome and translation machinery proteins

Untargeted analysis of our global quantitative proteomics dataset (Supplementary Dataset 5 and Supplementary Fig. 7) identified that ozonide and DHA treatment causes a significant dysregulation of 24 to 281 proteins within 3 h of drug exposure. Clustering analysis of all proteins significantly (P-value ≤ 0.05 and fold-change ≥ 1.5) perturbed by OZ277 treatment revealed that translation regulation (P-value = 5.184E^-7^) and the proteasome system (P-value = 5.288E^-4^) were the two main pathways affected. Parasite proteins in these two networks were significantly enriched following OZ277 treatment (Fig. 5). Similar protein clustering was also observed for DHA (Supplementary Fig. 11) treatment, with elevated levels of proteins involved in translation regulation (P-value < 1.0E^-9^) and the proteasome system (P-value 3.691E^-6^). A trend towards increased abundance of translation regulation and proteasome system proteins was also observed following 3 h of OZ439 exposure (Supplementary Dataset 5), however no protein networks were significantly enriched.

**Fig. 5.**
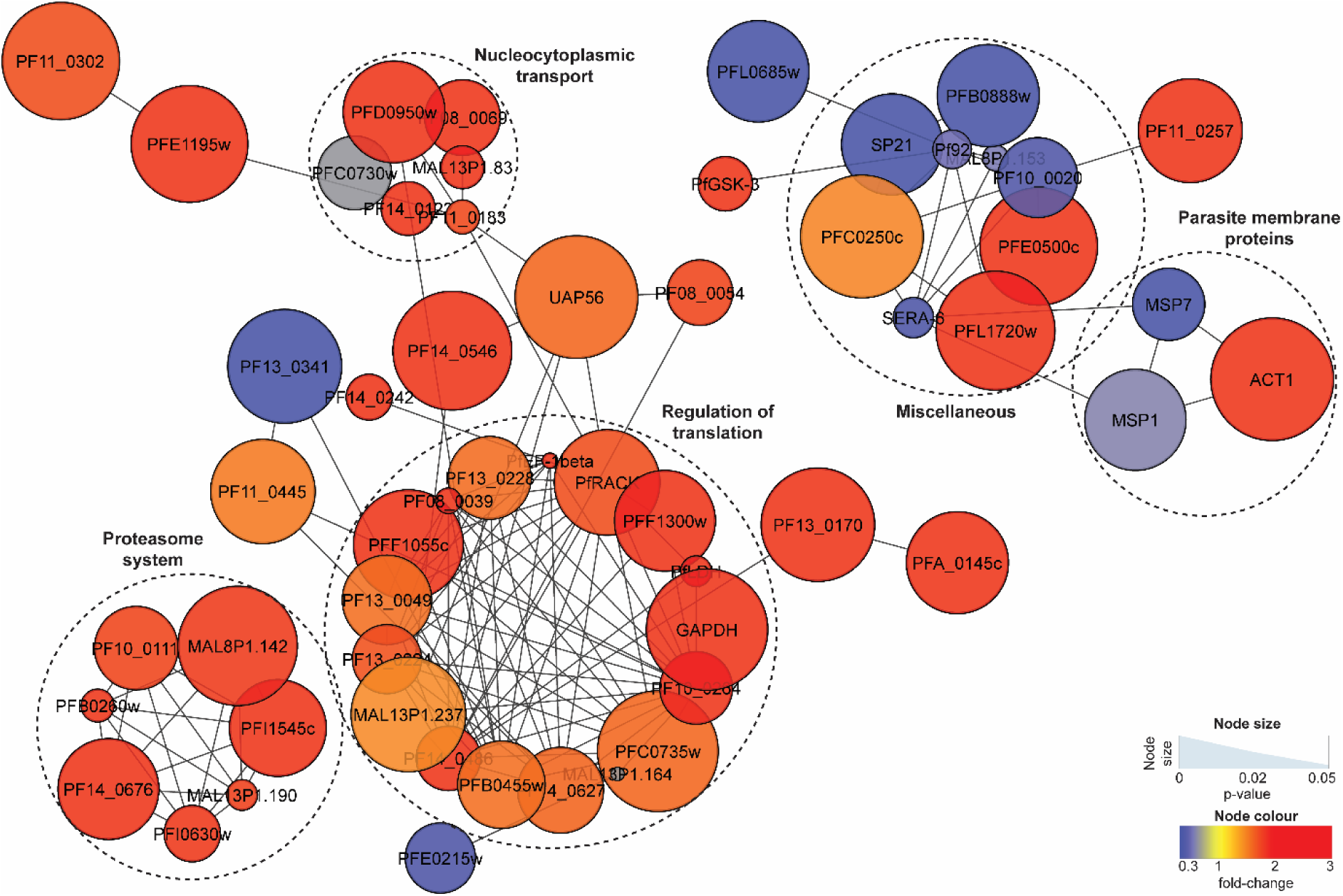
OZ277-induced disruption to the *P. falciparum* proteome. Network analysis of trophozoite stage parasite proteins perturbed following treatment with OZ277 (300 nM for 3 h). The network analysis was built using the STRINGdb interaction network analysis output (connectivity was based on experimental, database and co-expression evidence with a minimum interaction score of 0.7) in Cytoscape 3.6 with the ClusterONE algorithm. Node size represents P-value and node colour represents fold-change from at least three independent replicates.

### Extended exposure disrupts secondary metabolic pathways involved in ozonide antimalarial activity

Using untargeted metabolomic profiling, we also investigated whether parasite biochemical pathways other than Hb metabolism were affected following long-term ozonide treatment (> 3 h). Extended exposure induced metabolic perturbations beyond peptide metabolism, including, amino acid, lipid, cofactor and vitamin, and nucleotide metabolism (Fig. 6a and Supplementary Dataset 1). It is possible that some of these secondary responses to prolonged drug treatment represent nonspecific responses from dying parasites, but it is noted that different drug-specific responses were reported in similar metabolomics studies of other compounds [46–48], suggesting that these metabolic alterations are largely ozonide-specific. Extended treatment of trophozoite-stage parasites with OZ277 and OZ349 resulted in significant perturbations (P-value ≤ 0.05 and fold-change ≥ 1.5) to approximately 5% of the 217 putatively identified lipids (Supplementary Fig. 12). The major parasite neutral glycerolipid species, diglycerides (DG) and triglycerides (TG), were depleted within 6 h of ozonide exposure (Supplementary Fig. 12). DGs are the direct metabolic precursor of phosphatidylcholine (PC) and phosphatidylethanolamine (PE) lipids, the main glycerophospholipids in the parasite. Metabolites involved in PC and PE *de novo* synthesis accumulated in a time-dependent manner following extended ozonide treatment, while some of the PCs and PEs themselves, and other glycerophospholipids, were depleted (Fig. 6b and Supplementary Fig. 12). At the proteome level, four of the six enzymes in the *de novo* glycerophospholipid synthesis pathway (Kennedy pathway) were elevated after ozonide treatment compared to control (Fig. 6b and Supplementary Fig. 13). Prolonged treatment of rings (> 3 h) also disrupted *de novo* synthesis of PC and PE lipids (Supplementary Fig. 14).

**Fig. 6.**
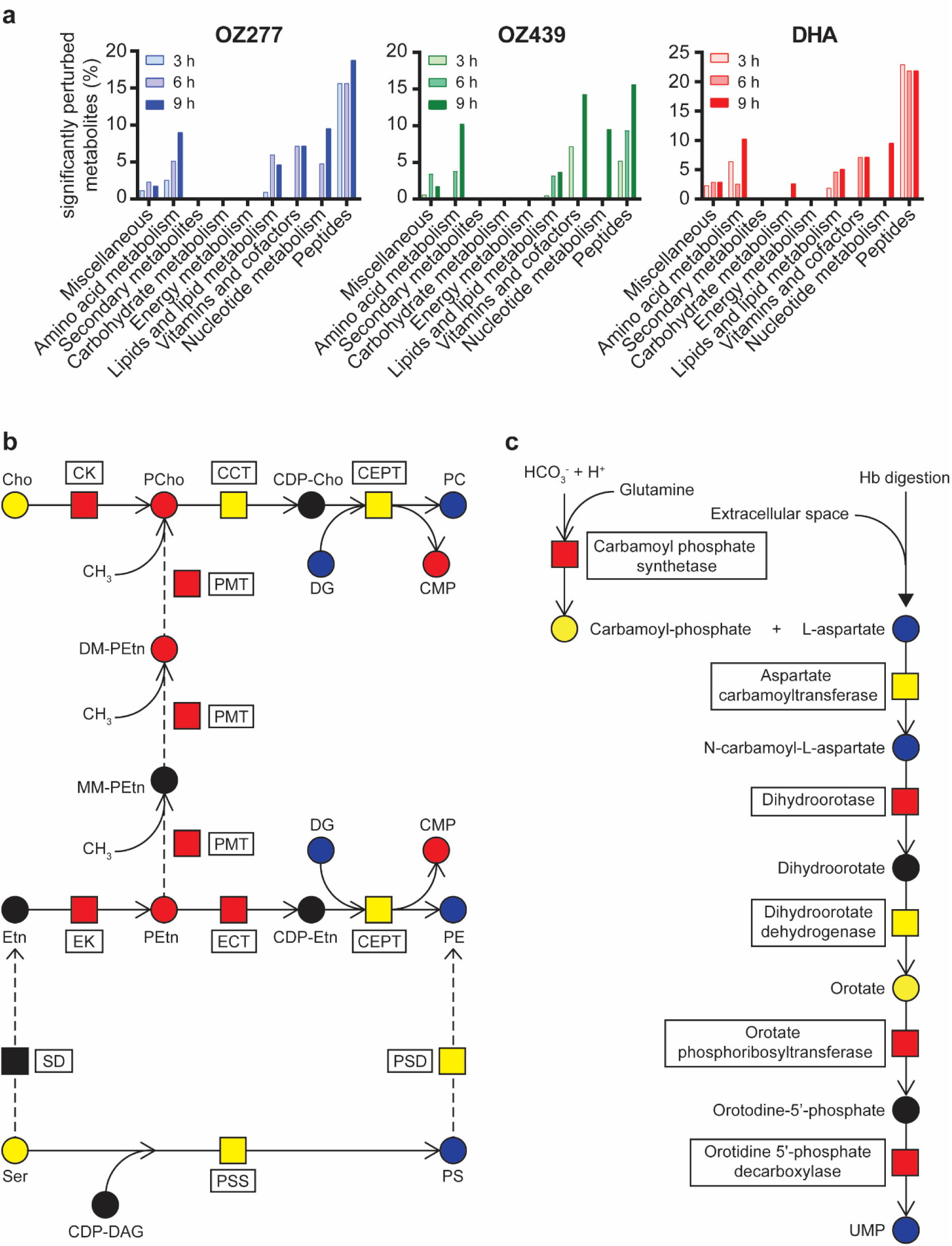
Peroxide-induced disruption of secondary biochemical pathways. **a**, Metabolic perturbations in trophozoite-stage parasite cultures. Pathway enrichment analysis showing the percentage of significantly perturbed metabolites (Welch’s *t* test; P*-*value < 0.05 and fold change > 1.5) as a function of metabolite class for extended (3, 6 and 9 h) exposure with OZ277, OZ439 (both 300 nM) and DHA (100 nM). **b**, Peroxide-induced disruption of the phosphatidylcholine (PC) and phosphatidylethanolamine (PE) lipid biosynthesis pathways within *P. falciparum* parasites. The dashed arrows represent an alternative route for the synthesis of PC from ethanolamine (Etn) in *P. falciparum*. CCT, choline-phosphate cytidyltransferase; CDP-, cytidine-diphospho-; CEPT, choline/ethanolamine phosphotransferase; Cho, choline; CK, choline kinase; CMP, cytidine monophosphate; DAG/DG, diglyceride; DM-, dimethyl-; ECT, ethanolamine-phosphate cytidyltransferase; EK, ethanolamine kinase; Etn, ethanolamine; MM-, monomethyl; PC, phosphatidylcholine; PCho, choline phosphate; PE, phosphatidylethanolamine; PEtn, ethanolamine phosphate; PG, phosphatidylglycerol; PMT, phosphoethanolamine N-methyltransferase; PS, phosphatidylserine; PSD, phosphatidylserine decarboxylase; PSS, phosphatidylserine synthase; SD, serine decarboxylase. **c**, Peroxide-induced perturbation of pyrimidine biosynthesis in trophozoite-stage parasite cultures. Hb, haemoglobin; UMP, uridine monophosphate. In **b**, **c**, metabolites (circles) and proteins (squares) in red and blue were increased and decreased in abundance after drug treatment, respectively. Yellow and black represent no change and not detected, respectively.

Extended ozonide exposure also disrupted parasite pyrimidine nucleotide biosynthesis. Metabolites of the *de novo* pyrimidine biosynthesis pathway, L-aspartate, N-carbamoyl-L-aspartate and uridine monophosphate (UMP), were all depleted after drug treatment (Fig. 6c and Supplementary Fig. 15), while at the proteome level, four of the six enzymes in this pathway were elevated compared to the control (Fig. 6c and Supplementary Fig. 16).

## Discussion

This study provides a detailed assessment of the *P. falciparum* biochemical pathways that are altered in response to ozonide antimalarial treatment. The “multi-omics” analysis revealed that ozonides act by rapidly perturbing parasite Hb catabolism prior to affecting other biochemical pathways, and suggested that the parasite regulates protein turnover to mitigate widespread ozonide-induced damage.

The time-resolved untargeted metabolomics approach allowed the mapping of primary and secondary ozonide-induced effects on parasite metabolism and revealed that short drug exposures induced rapid depletion of short-chain Hb-derived peptides, which was most pronounced in the more susceptible trophozoite-stage, in comparison to the less susceptible [28, 30] ring-stage parasites (Fig. 2 and Supplementary Fig. 3). It is noted that previous metabolomic profiling of ozonide-treated, magnetically purified *P. falciparum* cultures revealed no major alterations to parasite metabolism [48]. However, that was likely due to the high parasitaemia conditions (> 90%) of magnetically purified cultures inducing rapid ozonide degradation such that no measurable antimalarial activity could occur [28]. Consistent with previous reports [46, 47], our metabolomics analysis also showed that DHA induced depletion of short Hb-derived peptides. Interestingly, this occurred more rapidly for DHA than the ozonides (within 1.5 h of exposure versus 3 h of exposure), which agrees with the reported exposure time-dependence of activity for DHA and ozonides [30].

Digestion of host Hb is essential for parasite survival as it provides amino acids for parasite protein synthesis and has additional non-anabolic functions, such as maintaining osmotic stability of the iRBC [49–51]. Hb digestion is most active during the trophozoite-stage [52], resulting in extensive turnover of diverse Hb-derived peptides and the release of a high concentration of haem, which activates peroxide antimalarials within the parasite [3, 30, 53].

The extensive drug activation and Hb turnover in trophozoites likely explains the profound impact of peroxide antimalarials on Hb-derived small peptides in trophozoites (in terms of both the range of Hb peptides affected and magnitude of peptide depletion) compared to the ring-stage (Fig. 2). Although it is generally assumed that little Hb digestion occurs within ring-stage parasites, expression of active Hb-degrading proteases [30] and small haemozoin crystals (by-products of Hb digestion) have been detected [54–56]. This indicates that a low level of active Hb degradation occurs in ring-stage parasites and supports the observation of depleted Hb-derived small peptides following ozonide treatment (Fig. 2c and Supplementary Fig. 3 and 4).

The accumulation of longer Hb-derived peptides (Fig. 3b) and depletion of shorter di, tri and tetrapeptides (Fig. 2) suggests that the ozonides disrupt Hb catabolism through an inhibitory effect on the proteases involved in the conversion of large to small Hb peptides. However, we cannot rule out inhibition of the peptide transporters on the digestive vacuole membrane, or general impairment of digestive vacuole function, as contributors to this peptide phenotype. In response to perturbations within the digestive vacuole and impaired Hb catabolism, we propose that the parasite increases the abundance and activity of all proteases involved in Hb digestion, except falcilysin (Fig. 4a), which has been shown to localise to both the parasite apicoplast and the digestive vacuole, suggesting that it may have a function beyond Hb catabolism [60, 61]. It is noted that the activity of cysteine proteases involved in Hb digestion was elevated within 1 h of ozonide exposure, before the first time point of significant small peptide depletion (1.5 h), suggesting that the rapid peroxide effect on Hb catabolism occurs before there is a detectable change in small peptide levels using metabolomics.

Rapid disruption of Hb catabolism by the ozonides (and artemisinins) agrees with the hypothesis that peroxide-based drugs are activated by Fe(II) or haem to produce reactive intermediates in the parasite digestive vacuole where Hb digestion takes place. The resulting ozonide-derived radicals were recently shown to alkylate haem within iRBCs after short drug exposures [64] and the radicals may also alkylate and inactivate digestive vacuolar proteins, including the proteases involved in Hb digestion [19, 20, 27]. Our findings suggest that the digestive vacuole FPs are unlikely to be the Hb proteases initially targeted by the ozonides. The activity of these enzymes increased after ozonide treatment (Fig. 4b) and cysteine protease inhibition with E64d resulted in accumulation of full-length Hb and digestive vacuole swelling, whereas these features were not seen in peroxide treated-parasites (Supplementary Fig. 6). Ozonides (and artemisinins) are reported to alkylate proteins localised to the parasite digestive vacuole, including proteases involved in Hb digestion, for example, plasmepsins [19, 20, 27], which could be the initial intraparasitic protein targets of peroxide antimalarials. Consistent with our hypothesis that disrupted Hb catabolism is the key early event initiated by peroxide treatment, we demonstrated that the initial antiparasitic effects of peroxide exposure were enhanced when parasites were forced to rely solely on Hb degradation for amino acids (Fig. 4e). Taken together, we propose that Hb digestion is the initial pathway affected as a result of peroxide antimalarial treatment.

Depletion of short Hb-derived peptides following peroxide treatment was also confirmed in artemisinin resistant and sensitive parasite lines (Fig. 2d) that exhibit differential sensitivity to ozonides and DHA in short pulse assays [30, 31, 57]. The level of drug-induced peptide depletion was generally less extensive in the *K13*-mutant (Cam3.II^R539T^) compared to that in the drug-treated *K13*-wildtype revertant strain (Cam3.II^rev^) (Fig. 2d), suggesting that the peroxide impact on Hb catabolism is diminished in resistant parasites. This diminished affect may be a result of increased survival of *K13*-mutants following short peroxide exposure and could be mediated by altered Hb metabolism [38, 65, 66], augmented antioxidant defence pathways [38, 58, 59] or an enhanced stress response [32].

One caveat is that the precise origin of the depleted di, tri and tetrapeptides detected by untargeted metabolomic screening cannot be definitively determined due to their short sequences. MS/MS confirmation of the amino acid sequence was obtained for a subset of these depleted peptides and the confirmed sequences could be mapped to Hb. Furthermore, combined with additional lines of evidence pointing to a mechanism involving ozonide-induced disruption of Hb catabolism, it is likely that most of the small peptides that were perturbed by drug treatment originated from Hb. The short peptides unable to be mapped to Hb could originate from other RBC proteins or *Plasmodium* proteins, and could be associated with peroxide-induced inhibition of proteasome function and altered proteostasis [21]. It is important to note that the majority of peptides detected in the metabolome were not perturbed by drug treatment and most of these unaffected peptides could not be mapped to Hb. Furthermore, endogenous long peptides from only one *Plasmodium* protein were reproducibly perturbed by peroxide treatment. These data suggested that general parasite protein degradation was not significantly affected as a result of these short peroxide exposures.

Extended drug treatment (>3 h) induced disruption of additional biochemical pathways beyond Hb catabolism, including lipid and nucleotide metabolism (Fig. 6a), representing secondary biochemical pathways involved in peroxide activity. In both ring and trophozoite-stage parasites, drug treatment induced an accumulation of several metabolic intermediates in the *de novo* synthesis pathways of PC and PE lipids (known as the Kennedy Pathways) (Fig. 6b and Supplementary Fig. 12 and 14), which are the major lipid components of parasite membranes [46]. Proteomic analysis revealed that all enzymes directly upstream of the elevated metabolites in the Kennedy Pathways were also increased in abundance (Fig. 6b and Supplementary Fig. 13), possibly to increase the synthesis of PC and PE as a biochemical response to drug-induced membrane damage. These findings are consistent with the biological activity of ozonides and artemisinins involving non-specific damage to parasite membranes, such as the digestive vacuole and mitochondrial membranes [67, 68], through lipid peroxidation, which becomes apparent after an extended duration of drug exposure (> 3 h) and the production of reactive oxygen species [67, 69–72]. Inhibiting the synthesis of key phospholipids is detrimental to parasite survival [73] and it is likely that perturbation to this pathway contributes to peroxide antimalarial activity.

Prolonged ozonide and DHA exposure in trophozoite-stage parasites also led to the depletion of DGs and TGs (Fig. 6b and Supplementary Fig. 12). These are the two main neutral glycerolipid species in parasites [74] and these lipids increase in abundance as the asexual parasite matures, indicating their importance for growth and development [74]. DGs and TGs are packaged into neutral lipid bodies, which are closely associated with the parasite digestive vacuole [75–77]. Neutral lipid bodies are thought to concentrate free haem and catalyse its biocrystallisation into non-toxic haemozoin [76–78], placing them proximal to the location where peroxide antimalarials are thought to be activated. Furthermore, fluorescently-tagged artemisinin and ozonide derivatives have been shown to accumulate in neutral lipid bodies [70, 71]. Activated drug, or potentially alkylated haem adducts [64], may therefore promote oxidative damage to DGs and TGs within neutral lipid bodies and limit the availability of key neutral lipids that are required for parasite development [74]. As DGs are key precursors for membrane phospholipid synthesis (e.g. PC and PE), it is expected that DG depletion also contributes to the upregulation of *de novo* phospholipid biosynthesis (Kennedy) pathways.

Disruption of the parasite pyrimidine biosynthetic pathway at both the metabolite and protein levels (Fig. 6c and Supplementary Fig. 15 and 16) was also apparent in ozonide and DHA treated parasites. This finding is consistent with previous studies demonstrating DHA-induced alterations in parasite pyrimidine metabolism [46]. Furthermore, carbamoyl phosphate synthetase and aspartate carbamoyl transferase, which catalyse the initial steps of parasite pyrimidine biosynthesis, are reported to be alkylation targets of artemisinins, although this has not yet been shown for the ozonides [19]. Peroxide-induced inhibition of one or both of these initial pyrimidine biosynthetic enzymes may be responsible for the depletion of downstream pyrimidine biosynthesis intermediates, and a corresponding increase in the protein levels of some enzymes in this pathway as a compensatory response, as shown in our study.

The alternative hypothesis that these secondary pathways are non-specific responses in dying parasites is also possible. However, different drug-specific biochemical responses were reported in metabolomics, proteomics and peptidomics studies of other antimalarials, even when parasites were exposed to drugs for extended durations [46–48, 79, 80]. Combined with reports showing that peroxide antimalarials target proteins in both the phospholipid and pyrimidine biosynthesis pathways [19, 20, 27] and colocalise with neutral lipids within iRBCs [70, 71], the drug-specific biochemical responses of parasites detected in metabolomics studies suggests that the secondary metabolic alterations observed here are likely ozonide-specific and related to the pleiotropic effect of peroxides on parasite metabolism.

Our work demonstrating that peroxide antimalarials affect multiple aspects of parasite biochemistry is consistent with previous reports [46, 47]. Global proteomic analysis of ozonide and DHA-treated parasites revealed a pronounced enrichment of proteins involved in protein translation and the ubiquitin-proteasome system (Fig. 5 and Supplementary Fig. 11). Previous studies have shown that artemisinins inhibit protein translation and proteasome activity [21, 81], and the observed enrichment in these pathways from our study may represent a response to this inhibition, either by regulation of protein expression, or decreased degradation of these proteins. Furthermore, the enrichment of proteins in the translation and proteasomal pathways may reflect a general stress response to enhance protein turnover and mitigate peroxide-mediated cellular damage [82]. Peroxide-induced oxidative insult and widespread protein alkylation is thought to induce accumulation of damaged and misfolded proteins [22, 32, 83], and the parasite relies on translational regulation and a functional ubiquitin-proteasome system to restore proteostasis [84, 85].

Based on our findings, we have proposed a model for the MoA of peroxide antimalarials (Fig. 7). Hb-derived free haem activates the peroxide bond of ozonides (or artemisinins) within the digestive vacuole and the resulting drug-derived radicals initially alkylate haem [64] and damage proteases involved in Hb digestion leading to disruption of the Hb degradation pathway. In response to drug-induced damage, the parasite increases the abundance and activity of Hb-digesting proteases. Following prolonged exposure, drug-derived radicals induce further oxidative insult and cause widespread alkylation of parasite components, including damage to lipids, and proteins involved in other vital parasite functions. To mitigate drug-induced cellular damage, the parasite engages a proteostatic stress response involving upregulation of proteins involved in translational regulation and the ubiquitin-proteasome system. Parasite death ultimately occurs when drug-mediated damage overwhelms these parasite defensive mechanisms. In artemisinin resistance, *K13* mutations alter parasite Hb metabolism [38, 65, 66] and enhance antioxidant capacity [38, 58, 59] and stress response pathways [32], limiting the damage of drug-derived radicals and increasing parasite survival.

**Fig. 7.**
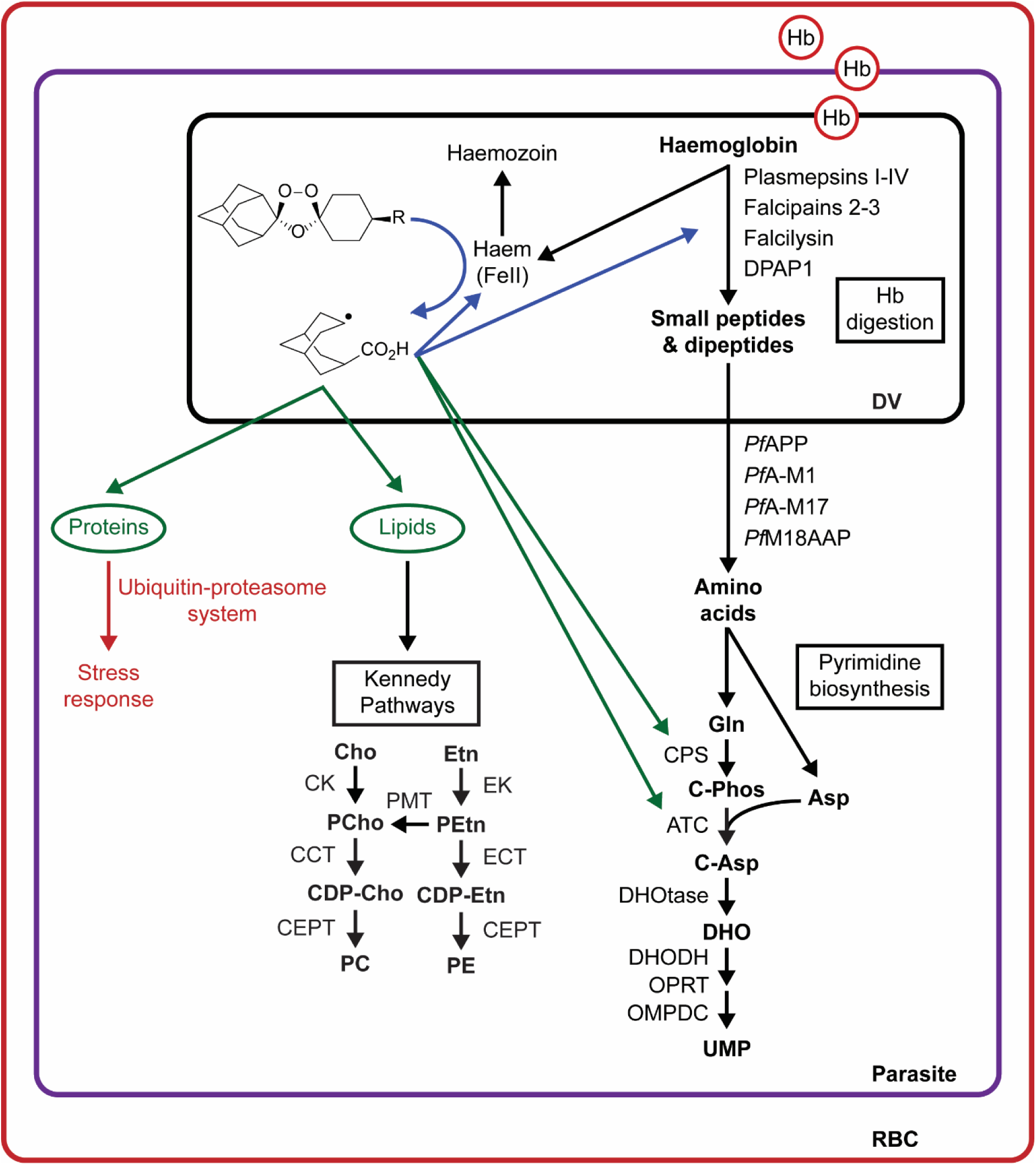
Proposed model for ozonide antimalarial activity in *P. falciparum* infected red blood cells. Haemoglobin-derived haem activates peroxide antimalarials within the parasite digestive vacuole. The resulting drug-derived radicals initially damage components proximal to the activation site, including proteases involved in haemoglobin digestion (arrow shown in blue). This leads to disruption of the haemoglobin degradation pathway. To correct for peroxide-induced damage, parasites may respond by increasing the abundance and activity of proteases involved in haemoglobin catabolism. Peroxide radicals induce further oxidative insult and cause widespread alkylation of parasite components as the duration of drug exposure is increased (arrows shown in green). This may include damage to lipids, inducing upregulation of the Kennedy Pathways, and proteins involved in other vital parasite functions, such as pyrimidine biosynthesis. To mitigate peroxide-induced cellular damage, the parasite engages a stress response involving translational regulation and the ubiquitin-proteasome systems. Asp, aspartic acid; ATC, aspartate carbamoyltransferase; C-Asp, carbamoyl-aspartate, C-Phos, carbamoyl-phosphate; CCT, choline-phosphate cytidyltransferase; CDP-, cytidine-diphospho-; CEPT, choline/ethanolamine phosphotransferase; Cho, choline; CK, choline kinase; CMP, cytidine monophosphate; CPS, carbamoyl phosphate synthetase; DHO, dihydroorotate; DHODH, dihydroorotate dehydrogenase; DHOtase, dihydroorotase; DV, digestive vacuole; ECT, ethanolamine-phosphate cytidyltransferase; EK, ethanolamine kinase; Etn, ethanolamine; Gln, glutamine; Hb, haemoglobin; OMPDC, orotodine 5’-phosphate decarboxylase; OPRT, orotate phosphoribosyltransferase; PC, phosphatidylcholine; PE, phosphatidylethanolamine; PCho, choline phosphate; PEtn, ethanolamine phosphate; PMT, phosphoethanolamine N-methyltransferase; RBC, red blood cell; UMP, uridine monophosphate.

Although DHA had a more rapid and pronounced effect on metabolism compared to the ozonides, these antimalarials affected similar biochemical pathways suggesting that they have a similar MoA. This could raise concerns for the deployment of ozonides in areas affected by artemisinin resistance [86], which is characterised by infections with parasites that can withstand the short DHA exposures observed in the pharmacokinetics of clinically-used artemisinins [17, 30]. However, the temporal metabolomics analysis demonstrated that prolonged peroxide exposure perturbed additional pathways beyond the digestive vacuole, raising the possibility that long half-life ozonides (e.g. OZ439) may impact additional parasite functions during prolonged exposure, and potentially overcome resistance associated with the short-lived artemisinins. There are mixed reports regarding ozonide activity in artemisinin resistant parasites, and further clinical studies are required to determine the potential utility of ozonides in artemisinin resistant malaria infections [64]. In the context of growing concerns about the spread of multi-drug resistant malaria parasites, this insight into the MoA of peroxide antimalarials, and the parasite’s response to treatment, offers potential avenues for targeting the malaria parasite with novel drug regimens that have improved antimalarial efficacy and limit the generation of drug resistance.

## Materials and Methods

### *Plasmodium falciparum* culture conditions

*P. falciparum* parasites (3D7 strain, Cam3.II^R539T^, and Cam3.II^rev^) were cultured as previously described [87]. RBCs were obtained from the Australian Red Cross Blood Service in Melbourne. Artemisinin resistant and sensitive *P. falciparum* isolates were kindly provided by Professor David Fidock, Columbia University and included the field-derived *K13*-mutant, Cam3.II^R539T^, and the *K13*-wildtype on an isogenic background (Cam3.II^rev^) [31]. Parasites were tightly synchronised by double treatment with sorbitol [88]. For 3D7 parasites, trophozoite or ring stage parasite cultures were adjusted to 10% parasitaemia and 2% Hct. For experiments using artemisinin resistant and sensitive isolates, trophozoite parasite cultures were adjusted to 4% parasiteamia and 2% Hct.

### Metabolomics sample preparation

In 3D7 parasites, comprehensive time-course analysis was performed to determine the peroxide-induced metabolic profile in ring- and trophozoite-stage parasites. Ring-stage parasite cultures (6-12 h post invasion) were exposed to drug (1 µM of OZ277 or OZ439, 300 nM of DHA and 0.03% DMSO) for 0, 3, 6 and 9 h prior to metabolite extraction, while trophozoite stage parasite cultures (28-34 h post invasion) were exposed to drug (300 nM of OZ277 or OZ439, 100 nM of DHA and < 0.03% DMSO) for 0, 0.5, 1.5, 3, 6 and 9 h.

In artemisinin resistant and sensitive isolates, prior to drug incubation, the age of paired artemisinin sensitive and resistant parasites was confirmed by analysis of Giemsa stained thin blood smears. Trophozoite parasite cultures (22-26 h) were exposed to 100 nM of DHA, OZ277 and OZ439 nM for a duration of 1, 3 and 5 h, respectively. Under the conditions used in this metabolomics study (4% parasitaemia and 2% Hct), these concentrations and exposure times were found to be sub-lethal in both the Cam3.II^R539T^ and Cam3.II^rev^ parasite lines (data not shown). Metabolomics studies included treatment of non-infected RBCs as controls and all experiments were performed on at least three independent occasions.

Following drug incubation, for ring-stage experiments (3D7), metabolites were extracted from 2 x 10^8^ cells using 200 µL of cold chloroform/methanol/water (1:3:1), while for trophozoite stage experiments (3D7), 1 x 10^8^ cells were used and metabolites extracted using 150 µL of cold chloroform/methanol/water (1:3:1). For trophozoite stage experiments (3D7) where long-term peroxide treatment was used (up to 9 h), metabolites were extracted from 1 x 10^8^ cells using 150 µL of cold methanol. For artemisinin resistant and sensitive isolates, 1 x 10^8^ cells were used and metabolites extracted using 150 µL of cold methanol. Metabolomics extraction was as previously described [47]. Insoluble precipitates were removed by centrifugation and 110 µL of metabolite extract was transferred to glass LC-MS vials and stored at −80 °C until analysis. A 15 µL aliquot of each sample was combined to generate a pooled biological quality control (PBQC) sample for analytical quality control and metabolite identification procedures.

### Metabolomics LC-MS analysis and data processing

Untargeted LC-MS analysis was performed using HILIC chromatography (ZIC-pHILIC; Merck®) with an alkaline mobile phase on an Ultimate U3000 LC system (Dionex) and Q Exactive Orbitrap MS (ThermoFisher®) operating in both positive and negative ion mode as previously described [38, 47]. The PBQC sample was run periodically throughout each LC-MS batch to monitor signal reproducibility and support downstream metabolite identification. Extraction solvent was used as blank samples to identify possible contaminating chemical species. To aid in metabolite identification, approximately 250 authentic metabolite standards were analysed prior to each LC-MS batch and their peaks and retention time manually checked using the ToxID software (Thermofisher®).

Metabolomics data were analysed using the IDEOM workflow [89]. High confidence metabolite identification (MSI level 1) was made by matching accurate mass and retention time to authentic metabolite standards [90]. Putative identifications (MSI level 2) for metabolites lacking standards were based on exact mass and predicted retention times [91]. Specifically, the identification of peptides was based on either accurate mass or a combination of accurate mass and MS/MS analysis, which allowed definitive confirmation of the amino acid sequence in selected peptides. In the ring and artemisinin resistance studies, LC-MS peak heights representing metabolite abundances were normalised by median peak height, while quality control procedures indicated that normalisation of metabolite abundances was not required in the 3D7 trophozoite studies. Univariate statistical analysis was performed using IDEOM and Welch’s *t* test [89]. Multivariate statistical analysis was performed on the mean centred and auto-scaled data using the web-based tools in Metaboanalyst [92]. Sparse partial least squares – discriminant analysis (sPLS-DA) algorithms were run with increasing numbers of metabolites in each component (up to 150 metabolites), with minor changes to the model when more than 10 metabolites were used. The final sPLS-DA plots shown in the supplementary data were developed using 10 metabolites in each component. Significant metabolites (P-value ≤ 0.05) were confirmed by manual integration of raw LC-MS peak areas in TraceFinder^TM^ (ThermoFisher®).

### Functional assays to measure haemoglobin abundance

The Hb fractionation assay was adapted from [35, 80]. Briefly, 3D7 trophozoite-stage parasites were incubated with DHA, OZ277, OZ439, E64d or a DMSO control for either 1 or 3 h. Following incubation, Hb, haem and haemozoin species were separated and measured using the Hb fractionation assay [35, 80], and smears were made using Giemsa stain to check for parasite viability and digestive vacuole morphology by light microscopy. For the Hb fractionation assay, the samples were normalised via a paired analysis to the DMSO control and graphed as their fold change vs DMSO ± SD. All fractions had >4 replicates from >2 independent experiments.

Hb monomer was also measured using SDS-PAGE gel. Briefly, 3D7 trophozoite-stage parasites were incubated with OZ277, E64d or a DMSO control for 3 h. Following incubation, extracted parasites were resolved on SDS-PAGE gels and proteins stained using Coomassie blue.

### Peptidomics sample preparation

Peptidomics samples were prepared as previously described with minor modifications [38]. Briefly, intracellular parasites were harvested after 3 h of drug treatment followed by trichloroacetic acid protein precipitation and centrifugal filtration using a 10 kDa cut-off filter (Amicon Ultra). The flow-through containing endogenous peptides was collected and peptide concentration was measured using a bicinchoninic acid (BCA) protein assay (Thermo Scientific Pierce) as per manufacturer’s protocol. Equal concentration of peptides (53-75 µg) were used for peptidomic analysis. Peptide samples were then subjected to desalting using in-house generated C18 StageTips [93]. The elutes were then dried and resuspended in 20 µL of 2% (v/v) acetonitrile (ACN) and 0.1% (v/v) formic acid (FA) for LC-MS/MS analysis.

### Peptidomics nanoLC-MS/MS analysis and data processing

LC-MS/MS analysis was performed using an Ultimate U3000 Nano LC system (Dionex) and Q Exactive Orbitrap MS (ThermoFisher®) as previously described [38].

Peptide identification was performed using PEAKS DB software [94]. Maximum mass deviation and false discovery rates were set at 0.5 Da and 0.01 respectively. No post translational modification or digestion were selected and identified peptide sequences were matched to *Homo sapiens* and *P. falciparum* databases. The mass to charge ratio and retention time of each identified peptide were imported into TraceFinder^TM^ (ThermoFisher®) and the peak intensity for each peptide was obtained manually. Further statistical analyses (Student’s *t* test) were performed using Microsoft Excel for paired drug treated and DMSO control samples.

### Proteomics sample preparation and triplex stable isotope dimethyl labelling

Proteomics samples were prepared as previously described with minor modifications [38]. Briefly, 1000 µg of protein, accurately determined using Pierce BCA assay, from each sample was incubated overnight with sequencing grade trypsin (Promega) (1:50) at 37℃. Quantitative triplex stable isotope dimethyl labelling was initiated on the following day using light, intermediate or heavy dimethyl labelling reagents [95]. The samples were then subjected to ion-exchange fractionation using a disposable Strong Cationic Exchange Solid Phase Extraction cartridge (Agilent Bond Elut) [96]. The fractions were desalted using in-house generated StageTips [93], dried and resuspended in 20 µL of 2% (v/v) ACN and 0.1% (v/v) FA for LC-MS/MS analysis.

### Proteomics nanoLC-MS/MS analysis and data processing

LC-MS/MS analysis was performed using an Ultimate U3000 Nano LC system (Dionex) and Q Exactive Orbitrap MS (ThermoFisher®) as previously described [38]. Protein identification and quantification was performed using the MaxQuant proteomics software [97]. The data analysis parameters in MaxQuant were set as previously described [38] with minor modifications. Dimethylation settings were adjusted to triplets; DimetNterm0 with DimetLys0, DimetNterm4 with DimetLys4 and DimetNterm8 with DimetLys8 were selected as light, intermediate and heavy label respectively. To correct for differences in protein amount between groups, the protein ratios were normalised in MaxQuant at the peptide level so that the log2 ratio is zero [97]. Known contaminants such as trypsin, keratin and reverse sequences were removed from the MaxQuant output. Fold-changes for the drug treated samples relative to the DMSO control samples were calculated in Microsoft Excel. One sample t-test was used to test the mean of combined experiment groups against the known mean (µ = 0) [98]. For each drug, the proteins that were identified in at least three independent experiments were filtered based on P-value (≤ 0.05) and fold-change (≥ 1.5) to generate a list of significantly perturbed proteins. The bioinformatics interaction network analysis tool STRINGdb [99] was used to build a protein-protein interaction network using the significantly perturbed proteins. Connectivity was based on experimental, database and co-expression evidence and a strict minimum interaction score (> 0.7) was applied to limit false positive associations in the predicted network. The STRINGdb protein connectivity output was exported to Cytoscape 3.6[100] and the ClusterONE algorithm was used to integrate and visualise relationships between proteins that were significantly perturbed by drug treatment.

### Temporal activity-based protein profiling of cysteine protease activity using activity-based probes

Activity-based probes (ABPs) were used to measure protease activity [43] following ozonide exposure according to established methods [101]. In these assays, tightly synchronised trophozoite stage parasites (28-34 h post invasion, 10% parasitaemia and 2% Hct) were treated with OZ277 (300 or 1000 nM) or OZ439 (300 nM) for up to 5 h. Untreated parasite controls contained an equivalent volume of DMSO (< 0.01%). Following ozonide treatment, parasites were purified by lysing the red blood cells using 0.1% saponin on ice. Parasite pellets were then lysed by sonication in citrate buffer (50 mM trisodium citrate [pH 5.5], 0.5% CHAPS, 0.1% Triton X-100, 4 mM dithiothreitol) or phosphate buffered saline (PBS) (pH 7.2). Supernatants were then cleared by centrifugation and transferred to new tubes. Protein concentration in each sample was determined using BCA protein assay (Pierce) and an equal amount of each sample was incubated with ABPs (DCG04; 2 µM or FY01; 1 µM) for 30 min at 37 °C to label active cysteine proteases. In experiments including the reversible cysteine protease inhibitor N-acetyl-Leu-Leu-Norleu-al (ALLN) (Merck), parasite lysates were pre-incubated with 10 µM of the inhibitor for 30 min prior to addition of the ABP for a further 15 min. In all cases, the reaction was quenched by the addition of 5x reducing buffer (50% glycerol, 250 mM Tris-Cl [pH 6.9], 10% SDS, 0.05% bromophenol blue, 6.25% beta-mercaptoethanol), boiled and separated by sodium dodecyl sulfate polyacrylamide (SDS-PAGE) on 15% polyacrylamide gels. As DCG04 is biotin-tagged, proteins were transferred to nitrocellulose membranes and incubated with streptavidin-AlexaFluor-647 followed by fluorescence detection with a Cy5 filter (excitation/emission: 649/670 nm) with an Amersham Typhoon 5 Biomolecular Imager (GE Healthcare Life Sciences). FY01 contains a Cy5 fluorophore, thus visualization of its targets was achieved by direct scanning of the gel for Cy5 fluorescence. Coomassie staining was used to confirm equal protein loading. Images were processed and quantified in either ImageJ 1.51f or Adobe Photoshop Creative Cloud 2017.

### Determination of antimalarial potency on parasites cultured in complete RPMI medium or medium lacking exogenous amino acids (except isoleucine)

Full RPMI medium 1640 (Sigma-Aldrich) contained all 20 amino acids and was supplemented with 5.94 g/l HEPES, 2.1 g/l NaHCO_3_, 50 mg/l hypoxanthine and 5 g/l Albumax II (Lifetech), making AA medium. RPMI medium 1640 lacking all amino acids (Life Research) was supplemented with isoleucine (Sigma-Aldrich) at a final concentration of 147.5 µM to make Iso medium [44]. All other supplements (HEPES, NaHCO_3_, hypoxanthine and Albumax II) were added as for AA medium above. To examine parasite susceptibility to the peroxide antimalarials in the Iso and AA mediums, drug pulse activity assays were performed as previously described [102]. Briefly, the medium of iRBC cultures containing 30 h trophozoite-stage parasites (10% parasitaemia and 2% Hct) was replaced with either Iso or AA medium immediately before initiating drug treatment. Parasites were treated with 300 nM of pyrimethamine, OZ277 or OZ439 or 100 nM of DHA for 3 h. Following the incubation period, drugs were removed by washing the cultures as previously described [28] using either AA or Iso medium supplemented with 2-5% Albumax II. Cultures were then adjusted to 0.5% parasitaemia and 2% Hct (final volume, 200 µL) as previously described [28] with either Iso or AA medium. After 48 h the parasitaemia was measured by counting Giemsa stained thin blood smears. Parasite survival was determined by graphing the parasitaemia for each test compound relative to the appropriate untreated medium (Iso or AA) control, which was set to 100%. All assays were performed in triplicate in at least three independent experiments.

## Data availability

Proteomics and Peptidomics mass spectrometry data and search results have been deposited to the ProteomeXchange Consortium via the PRIDE [103] partner repository with the dataset identifier PXD014313 and project name ‘System-wide biochemical analysis reveals ozonide and artemisinin antimalarials initially act by disrupting malaria parasite haemoglobin digestion’. Reviewers can access the dataset by using the username ‘’ and password ‘RQivOi19’.

Metabolomics spectrometry data is available at the NIH Common Fund’s Metabolomics Data Repository and Coordinating Center website, the Metabolomics Workbench http://metabolomicsworkbench.org, where it has been assigned Project ID PR000809. The data can be accessed directly via its project DOI: 10.21228/M83X38.

## Acknowledgments

The authors thank Professor Matthew Bogyo (Stanford University School of Medicine) and Doctor Edgar Deu (The Francis Crick Institute) for providing the FY01 probe. Doctor Edgar Deu is also gratefully acknowledged for advice regarding activity-based protein profiling experiments. The authors thank Professor David Fidock (Columbia University) for provision of the genetically modified Cam3.II^R539T^ and Cam3.II^rev^ *P. falciparum* isolates. The Monash Proteomics and Metabolomics Facility (Parkville Node) provided technical assistance with metabolomics and proteomics experiments. The Australian Red Cross Blood Service in Melbourne donated human red blood cells for *in vitro* parasite cultivation. Funding support was provided by the Australian National Health and Medical Research Council (NHMRC) project grants #APP1128003 and #APP1160705 and fellowship to D.J.C. (#APP1148700). L.E.-M is funded by an Australian Research Council (ARC) Discovery Early Career Researcher Award (DECRA) Fellowship (DE180100418) and the Grimwade Fellowship funded by the Russell and Mab Grimwade Miegunyah Fund at the University of Melbourne.

## Author contributions

C.G., G.S., S.A.C., and D.J.C. designed the experiments. C.G., G.S., A.D.P., and B.M.A executed the experiments. C.G., and G.S performed the mass spectrometry experiments. C.G., and G.S. analysed the data. C.G., G.S., L.E.-M., S.A.C., and D.J.C. wrote the manuscript. D.J.C supervised the study.

## Supporting information

Supplementary Dataset 1. IDEOM metabolomics output for OZ277, OZ439 and DHA treatment of *P. falciparum* (3D7 strain) ring and trophozoite infected RBC cultures and uninfected RBC cultures.

Supplementary Dataset 2. IDEOM metabolomics output for OZ277, OZ439 and DHA treatment of Cam3.II^R539T^ (artemisinin resistant) and Cam3.II^rev^ (artemisinin sensitive) *P. falciparum* parasite lines.

Supplementary Dataset 3. Peptidomics dataset for OZ277, OZ439 and DHA treatment (3 h) of *P. falciparum* trophozoite infected RBC cultures (3D7 strain).

Supplementary Dataset 4. Peptidomics dataset for DHA (1 h) treatment of *P. falciparum* trophozoite infected RBC cultures (Cam3.II^rev^, artemisinin sensitive).

Supplementary Dataset 5. Proteomics dataset for OZ277, OZ439 and DHA treatment of *P. falciparum* trophozoite infected RBC cultures (3D7 strain).

